# Mitonuclear interactions influence Alzheimer’s disease risk

**DOI:** 10.1101/654400

**Authors:** Shea J Andrews, Brian Fulton-Howard, Christopher Patterson, G Peggy McFall, Alden Gross, Elias K Michaelis, Alison Goate, Russell H Swerdlow, Judy Pa, for the Alzheimer’s Disease Neuroimaging Initiative

## Abstract

We examined the associations between mitochondrial DNA haplogroups (MT-hg) and their interactions with a polygenic risk score based on nuclear-encoded mitochondrial genes (nMT-PRS) with risk of dementia and age of onset of dementia (AOO). Logistic regression was used to determine the effect of MT-hgs and nMT-PRS on dementia at baseline (332 controls / 204 cases). Cox proportional hazards models were used to model dementia AOO (n=1047; 433 incident cases). Additionally, we tested for interactions between MT-hg and nMT-PRS in the logistic and Cox models. MT-hg K and a one SD larger nMT-PRS were associated with elevated odds of dementia. Significant antagonistic interactions between the nMT-PRS and MT-hg K and T were observed. Individual MT-hg were not associated with AOO; however, a significant antagonistic interactions was observed between the nMT-PRS and MT-hg T and a synergistic interaction between the nMT-PRS and MT-hg V. These results suggest that MT-hgs influence dementia risk, and that variants in the nuclear and mitochondrial genome interact to influence the age of onset of dementia.

**Highlights:**

- Mitochondrial dysfunction has been proposed to influence dementia risk
- MT-hg K and T interacted with a genetic risk score to reduce dementia risk
- MT-hg T and V interacted with a genetic risk score to influence dementia age of onset

## 1. Introduction

Alzheimer’s disease (AD) is a progressive neurodegenerative disease characterized by cognitive and functional deterioration resulting in a loss of independent living, and ultimately death (Masters et al., 2015). The neuropathological hallmarks of AD are the abnormal aggregation and accumulation of amyloid-β peptides into extracellular amyloid plaques and hyperphosphorylated tau intracellular neurofibrillary tangles, accompanied by neuroinflammation, gliosis, and neurodegeneration (Masters et al., 2015; Mhatre et al., 2015). As such, studies on AD pathogenesis and therapeutics have largely focused on the role of Aβ and Tau. However, with several negative trials of drugs targeting Aβ pathways, there has been increasing interest in evaluating the role of other pathological features in AD, such as mitochondrial dysfunction (Panza et al., 2019; Perez Ortiz and Swerdlow, 2019).

Mitochondria are vital to cellular function, firstly as the major source of cellular energy through the generation of adenosine triphosphate (ATP) via oxidative phosphorylation, but also through regulation of calcium uptake, apoptosis, and production and sequestration of reactive oxygen species (Gorman et al., 2016). Each mitochondrion possesses its own 16,569 base pair circular genome (mtDNA) that encodes 37 genes: 13 protein-coding genes, 22 tRNAs, and two ribosomal RNAs (Taanman, 1999). Genetic variation in the mitochondria is often described by established haplotype groups defined by a specific combination of single nucleotide polymorphisms (SNPs) that represent major branch points in the mitochondrial phylogenetic tree (van Oven and Kayser, 2009). The nuclear genome also plays a key role in mitochondrial function as it contains 1,145 genes that encode proteins that influence mitochondrial function (mitonuclear genes) (Calvo et al., 2016). These Mitonuclear genes encode the majority of the proteins involved in the oxidative phosphorylation system and are also essential for maintaining mtDNA replication, organelle network proliferation and destruction (Chinnery and Hudson, 2013). A recent systematic review of 43 studies examining the effects of mitonuclear incompatibility across vertebrates and invertebrates finding significant effects on health, including gene expression, metabolic traits, anatomical or morphological traits, lifespan and fecundity (Dobler et al., 2018)reported incompatibility between nuclear and mitochondrial genes can influence biological function. Furthermore, in six admixed human populations, increasing discordance between nuclear and mtDNA ancestry was associated with reduced mtDNA copy number - a proxy measure for mitochondrial function (Zaidi and Makova, 2019). Thus mitochondrial function relies on fine-tuned mitonuclear interactions that require the nuclear and mitochondrial genomes to be compatible with one another.

The central nervous system is particularly vulnerable to impaired mitochondrial metabolism because of its high-energy demands. Increasing evidence links mitochondrial dysfunction to neurodegenerative diseases such as AD. Support for the role of mitochondria in AD comes from studies observing changes in the rate of metabolism, disruption of fusion and fission, altered concentration of mitochondria in CSF, morphological changes and aggregation of Aβ in the mitochondria (Perez Ortiz and Swerdlow, 2019; Swerdlow, 2018). Additionally, maternal history of AD confers an increased risk of AD, cognitive aging and elevated biomarkers for AD, which is consistent with the maternal inheritance of mtDNA (Honea et al., 2012; Swerdlow, 2018). Despite this evidence, the role of the mitochondrial genome in AD remains inconclusive, as a recent systematic literature review of 17 studies reported few definitive findings on the association of mitochondrial genetic variation with AD (Ridge and Kauwe, 2018). In addition, while candidate gene studies have implicated several mitonuclear genes in AD risk, genome-wide association studies (GWAS) have not supported the association of specific mitonuclear genes with AD, with the exception of *TOMM40* which is in high linkage disequilibrium with *APOE* (Chiba-Falek et al., 2018; Kunkle et al., 2019). To date, no study has investigated whether the genetic variation in mitonuclear genes interacts with the mitochondrial genome to influence AD risk.

In this study, we investigate the association of mitonuclear interactions in Alzheimer’s disease by evaluating the interactions between an AD polygenic risk score that included only variants from mitonuclear genes (nMT-PRS) and mitochondrial haplogroups (MT-hg) on Alzheimer’s disease risk and survival.

## 2. Methods

### 2.1 Alzheimer’s Disease Neuroimaging Initiative

Data used in the preparation of this article were obtained from the Alzheimer’s Disease Neuroimaging Initiative (ADNI) database (adni.loni.usc.edu). The ADNI was launched in 2003 as a public-private partnership, led by Principal Investigator Michael W. Weiner, MD. The primary goal of ADNI has been to test whether serial magnetic resonance imaging (MRI), positron emission tomography (PET), other biological markers, and clinical and neuropsychological assessment can be combined to measure the progression of mild cognitive impairment (MCI) and early Alzheimer’s disease (AD).

Descriptive characteristics of ADNI participants at baseline and last assessment are presented in Table 1.

**Table 1:** Descriptive characteristics of ADNI at baseline and last assessment (survival analysis).

| Variable | Baseline Analysis |  | Survival Analysis |  |  |
| --- | --- | --- | --- | --- | --- |
|  | Controls<br>(n = 301) | AD<br>(n = 187) | Controls<br>(n = 273) | MCI<br>(n = 350) | AD<br>(n = 424) |
| Age, $\mu$ (sd) | 75.49 (5.23) | 75.67 (7.92) | 78.82 (6.78) | 77.83 (7.7) | 74.77 (8.14) |
| Gender, n (%) | 135 (44.85) | 82 (43.85) | 126 (46.15) | 134 (38.29) | 172 (40.57) |
| Education, $\mu$ (sd) | 16.3 (2.72) | 14.96 (2.98) | 16.44 (2.72) | 15.84<br>(2.82) | 15.42 (2.91) |
| APOE, n (%) |  |  |  |  |  |
| $\epsilon 3/\epsilon 3$ | 201 (60.54) | 60 (29.41) | 172 (63) | 183 (52.29) | 140 (33.02) |
| $\epsilon 4+$ | 92 (27.71) | 139 (68.14) | 69 (25.27) | 139 (39.71) | 278 (65.57) |
| $\epsilon 2+$ | 39 (11.75) | 5 (2.45) | 32 (11.72) | 28 (8) | 6 (1.42) |
| nMT-PRS, $\mu$ (sd) | -0.37 (0.85) | 0.47 (0.93) | -0.38 (0.86) | -0.08 (0.82) | 0.4 (0.94) |
| Haplogroups, n (%) |  |  |  |  |  |
| H | 169 (50.9) | 92 (45.1) | 130 (47.62) | 151 (43.14) | 200 (47.17) |
| I | 14 (4.22) | 6 (2.94) | 10 (3.66) | 10 (2.86) | 17 (4.01) |
| J | 29 (8.73) | 20 (9.8) | 25 (9.16) | 36 (10.29) | 38 (8.96) |
| K | 34 (10.24) | 31 (15.2) | 28 (10.26) | 46 (13.14) | 43 (10.14) |
| T | 31 (9.34) | 16 (7.84) | 23 (8.42) | 40 (11.43) | 41 (9.67) |
| U | 35 (10.54) | 26 (12.75) | 35 (12.82) | 48 (13.71) | 51 (12.03) |
| V | 11 (3.31) | 7 (3.43) | 13 (4.76) | 10 (2.86) | 14 (3.3) |
| W | 3 (0.9) | 4 (1.96) | 3 (1.1) | 7 (2) | 10 (2.36) |
| X | 6 (1.81) | 2 (0.98) | 6 (2.2) | 2 (0.57) | 10 (2.36) |

### 2.2 Nuclear DNA

GWAS data for ADNI participants were obtained and processed as previously described (Saykin et al., 2015). Briefly, genomic DNA extracted from blood were genotyped on Illumina GWAS arrays (ADNI1: 610-Quad; ADNI GO/2 OmniExpress). Genotype data then underwent stringent quality control checks, with variants excluded if the call rate was < 0.95, MAF was < 1% or were not in Hardy-Weinberg equilibrium (p < 1 × 10−6) and samples excluded if call rate was < 0.95, discordant sex was reported, cryptic relatedness, non-European ancestry or outlying heterozygosity. To empirically determine ancestry, the samples were projected onto principal components from known ancestral populations in the 1000 Genomes Project, with samples determined to be European population outliers if they were ± 6 SD away from the EUR population mean on the first 10 principal components (1000 Genomes Project Consortium et al., 2015). Within-ancestry principal components were created using the --PCA function in PLINK (Purcell et al., 2007) to correct for residual population stratification within the European population subset. SNPs that were not directly assayed were imputed using the Haplotype Reference Consortium (McCarthy et al., 2016), with imputed variants excluded due to poor imputation quality (INFO < 0.3) or low MAF (< 1%).

### 2.3 Mitochondrial DNA

138 mtDNA variants were available for 757 samples from ADNI1 who were genotyped on the Illumina 610-Quad array. Additional mitochondrial genetic variants were made available via imputation of the mitochondrial genome, as previously described (McInerney et al., 2019), using a custom reference panel of mitochondrial genome sequences and the chromosome X imputation protocol in IMPUTE2 (Howie et al., 2009). An additional 809 samples with mitochondrial variants were made available via whole genome sequencing (Ridge et al., 2018). MT-hg were assigned to the genotyped/imputed dataset (SNPs with an info score > 0.4) using Haplogrep2 (Weissensteiner et al., 2016), while in the whole genome sequenced dataset MT-hg were assigned using Phy-Mer (Navarro-Gomez et al., 2015).

### 2.4 Polygenic Risk scores

The software package PRSice was used to construct an AD PRS for nuclear encoded mitochondrial polygenic risk scores (nMT-PRS) (Euesden et al., 2015). To generate a mitonuclear AD PRS, SNPs from stage 1 of the International Genomics of Alzheimer’s Project (IGAP) (Lambert et al., 2013) were annotated to known protein-coding genes (±50kb) using MAGMA (de Leeuw et al., 2015) and those SNPs that were assigned to any of 1,158 mitonuclear genes were extracted (Calvo et al., 2016). A p-value threshold of 0.5 was used for inclusion of SNPs into the nMT-PRS as this threshold has been previously shown to have the most significant association with case/control diagnosis (Escott-Price et al., 2015). To obtain independent loci, LD clumping was performed by excluding SNPs that had an r2 > 0.1 with another variant with smaller p-value association within a 250kb window. SNPs were weighted by their effect sizes in IGAP. A total of 12,838 SNPs were included in the nMT-PRS.

### 2.5 Statistical Analysis

#### Cross-sectional analysis

The effect of the MT-hgs on baseline risk of dementia was assessed using Binomial multivariate logistic regression models with MT-hg H used as the reference group and adjusting for age, *APOE* status, sex, and the first two principal components. Multiplicative and additive models of interaction were used to examine the interaction between MT-hgs and the nMT-PRS. To test for multiplicative interactions between MT-hgs and nMT-PRS, an interaction term for the two variables was added in the model. Departures from additivity were tested using the relative excess risk due to interaction (RERI), which can be interpreted as the risk that is additional to the risk that is expected on the basis of the addition of the OR of the variables. In the absence of an interaction effect, the RERI is equal to 0. As MCI is an unstable diagnosis with individuals either converting to dementia, remaining stable or reverting back to normal cognition (Canevelli et al., 2016), participants with MCI were excluded from the analysis.

#### Survival analysis

In the survival analysis, age was used as the time to event scale. For subjects who were cognitively normal or suffering from MCI at baseline, AD age at onset was used. For participants with AD at baseline, age at onset was estimated. A Cox-proportional hazards model with adjustment for *APOE* status, sex, and the first two principal components was used to assess the effects of the nMT-PRS and MT-hgs on Alzheimer's age of onset. To evaluate potential interactions between the MT-hgs and the nMT-PRS, an interaction term was included in the model.

#### Sensitivity analysis

As a sensitivity analysis, an additional polygenic score was constructed (PRS w/o nMT & APOE) composed of all SNPs associated with late onset AD at p < 0.5 in IGAP, except for those annotated to known mitonuclear genes and the *APOE* region (±250kb of *APOE*). A total of 191,990 SNPs were included in the PRS. The cross-sectional and survival analysis were repeated introducing this additional PRS as a covariate.

All analysis was performed in the R 3.5.2 statistical computing environment.

## 3. Results

### 3.1 Association of mitonuclear interactions with Alzheimer’s risk

Binomial logistic regression was used to evaluate the main effects of the MT-hgs and nMT-PRS and their interaction on the likelihood of participants having AD (Table 2). In the main effects model, MT-hg K was associated with an increased risk of developing Alzheimer’s disease (OR: 2.03 [95% CI: 1.04, 3.97]), while MT-hg U was nominally associated with increased risk (OR: 1.99 [95% CI: 0.99, 3.97]). A 1 SD increase in the nMT-PRS was associated with an increased likelihood of developing AD (OR: 2.2 [95% CI: 1.68, 2.86]).

**Table 2:**
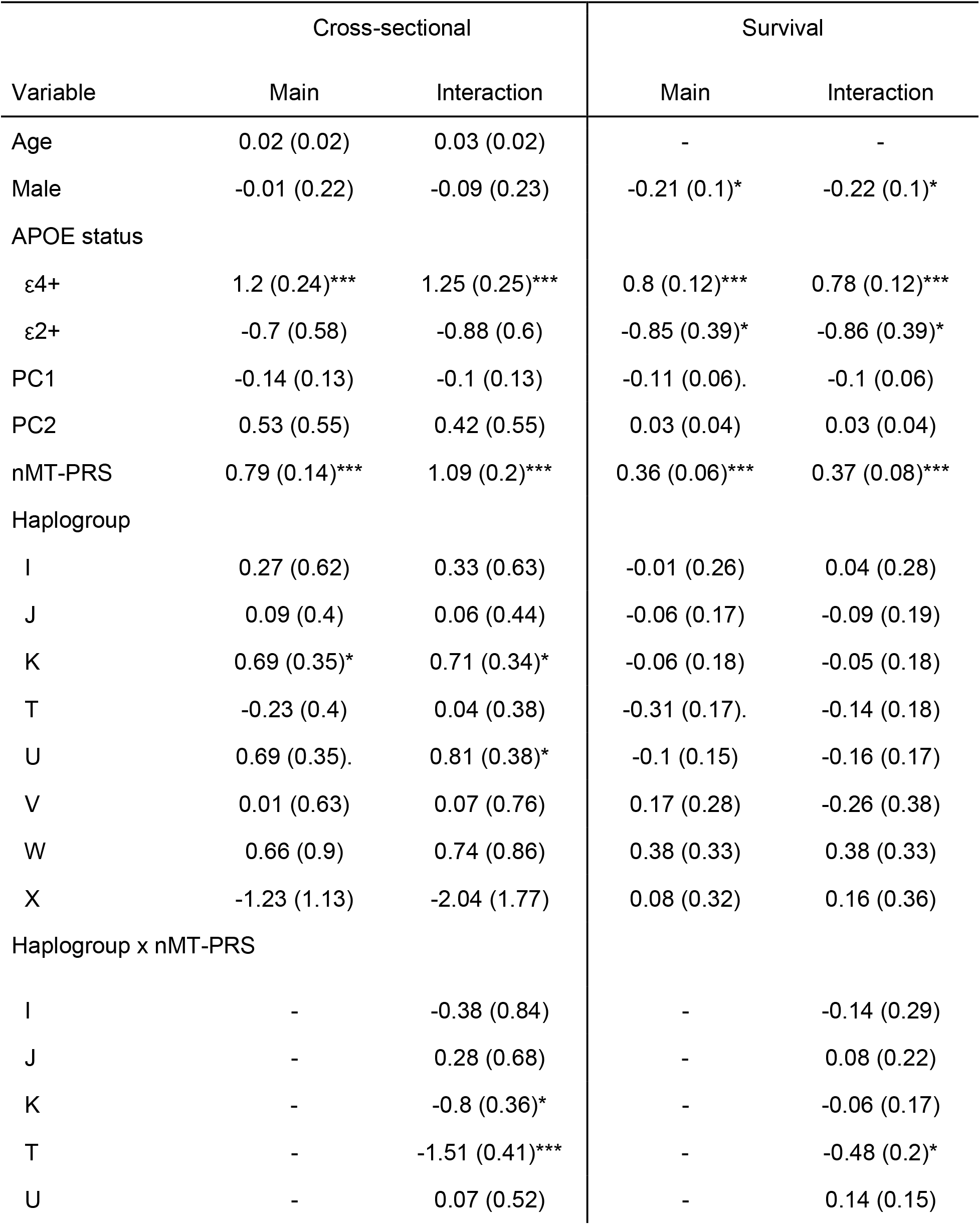
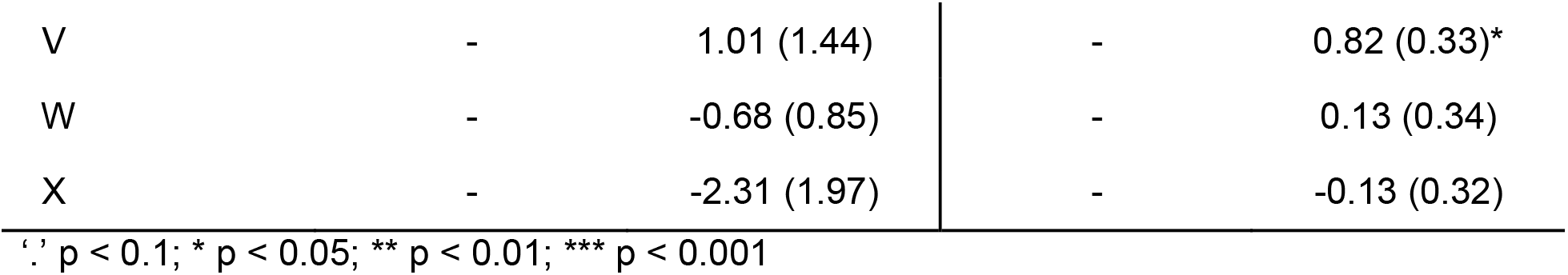
Association of mitonuclear interactions with baseline risk of Alzheimer's disease and Alzheimer’s age of onset (β [SE]).

In the interaction model, a significant interaction was observed between the nMT-PRS and MT-hg T (OR: 0.22 [95% CI: 0.1, 0.49]) and K (OR: 0.45 [95% CI: 0.22, 0.9]). Under an additive model, the RERI for MT-hg T and K were RERI −2.8 (95% CI: −4.33, −1.26) and −3.56 (95% CI: −5.54, −1.57) respectively, indicating that the nMT-PRS and MT-hg K and T acted antagonistically in relation to AD risk, such that the relative risk of AD was 2.8 and 3.56 times lower than expected from the addition of the separate effects of the nMT-PRS and MT-hg.

### 3.2 Association of mitonuclear interactions with Alzheimer’s age of onset

A Cox-proportional hazard model was used to evaluate the main effects of the MT-hgs and nMT-PRS and their interaction on Alzheimer’s age of onset (AOO) (Table 2). In the main effects model, a 1 SD increase in the nMT-PRS was associated with an earlier AOO (HR: 1.44 [95% CI: 1.28, 1.61]). However, none of the MT-hgs were significantly associated with AOO.

In the interactive model, a significant interaction was observed between the nMT-PRS and MT-hg T (HR: 0.62 [95% CI: 0.42, 0.91]) and V (HR: 2.28 [95% CI: 1.19, 4.35]). Under an additive model, RERI for MT-hg T and V were −0.7 (95% CI: −1.24, −0.16) and 1.06 (95% CI: −0.9, 3.02) respectively, indicating that the relative risk of AD was 0.7 times lower for MT-hg T and 1.06 times higher for MT-hg V then expected from the addition of the separate effects of the nMT-PRS and MT-hg.

### 3.3 Sensitivity analysis

The effect of adjusting the baseline logistic model and the survival model with a PRS risk score composed of non-nuclear mitochondrial SNPs and non-APOE region SNPs are presented in Supplementary Table 1. The effect of the nMT-PRS on AD risk and AOO was attenuated but remained statistically significant. The significant interactions observed between the nMT-PRS and MT-hg T in the baseline model and MT-hg V remained statistically significant after covarying for the PRS w/o nMT & APOE. However, the interaction with MT-hg K in the baseline model was nominally significant, while the interaction with MT-hg T was no longer significant in the survival model.

## 4. Discussion

In this study, we investigated whether mitonuclear interactions influence Alzheimer’s risk and survival by evaluating the interactive effects of MT-hgs and a polygenic risk score composed of mitonuclear genes (nMT-PRS) on baseline risk of AD and age of onset of dementia (AOO). We observed that nMT-PRS was associated with an increased risk of AD and an earlier AOO, even after adjusting for a PRS composed of SNPs from the rest of the genome. Additionally, we observed that MT-hg K was associated with an increased baseline risk of Alzheimer’s disease and in the interactive model, modified the risk associated with the nMT-PRS in an antagonistic manner, such that the combined effect of the nMT-PGS and MT-hg K was smaller than expected given their additive effects. In effect they are less harmful together then they are on their own, though the potential underlying mechanisms of this compensatory effect are complex (Lehner, 2011). MT-hg T was also observed to modify the risk associated with the nMT-PRS in an antagonistic manner for both baseline risk of AD and AOO. Finally, we observed that MT-hg V was associated with an increased risk of AD beyond that expected in the additive model with the nMT-PRS.

These results suggest that epistasis between nuclear and mitochondrial genomes, in which one gene’s effect is dependent on the presence of another gene or set of genes, influences the risk of Alzheimer’s disease. While to date, no previous study to our knowledge has investigated the interaction between mitonuclear genes and the mitochondrial genome in the context of AD, several studies have investigated associations between mitochondrial genetic variation and *APOE*. In *APOE* ε4 carriers MT-hg K and U were observed to have neutralizing effect (Carrieri et al., 2001; Maruszak et al., 2011) on AD risk. Conversely, SNP mt7028C, a defining SNP for MT-hg H, and MT-hg H5a had acted synergistically with *APOE* ε4 to increased risk of AD (Coto et al., 2011; Maruszak et al., 2011). Finally, SNP mt4336C which defines MT-hg H5a was associated with an increased risk of AD only in *APOE* ε4 carriers (Edland et al., 2002). Outside of Alzheimer’s disease, mitonuclear interactions have also been implicated in altering the penetrance of primary pathological mutations underlying mitochondrial disease or modifying the pathogenic phenotype of other diseases, such as non-syndromic sensorineural deafness (Kokotas et al., 2007; Morrow and Camus, 2017). The results of this study, in addition to the prevalence of mitonuclear epistasis in other diseases, suggests that the inconclusive results of mitochondrial genetic variation in AD may not only be due to small sample sizes, limited genetic data collection and inadequate approaches to association analysis (Ridge and Kauwe, 2018), but could also be attributed to the modifying effects of nuclear-encoded mitochondrial genes. As such, future studies investigating the association of mtDNA with AD should consider evaluating the modifying effect of nDNA.

The MT-hg association analysis is in agreement with two previous studies conducted in ADNI. Lakatos et al (Lakatos et al., 2010) investigated the association of four MT-hg clusters (HV, JT, UK, and IWX) with AD in 358 participants and found that the UK MT-hg cluster was associated with an increased risk of AD. Ridge et al (Ridge et al., 2013) utilized a TreeScanning approach to assess the relationship of mitochondrial genetic variation with structural MRI and cognitive biomarkers and found SNPs defining either MT-hg K1A1B or K1A1B2A1 and MT-hg U5B1 or U5B1B2 were associated with reduced temporal pole thickness, which is considered evidence of increased risk for AD. However, within the context of the wider literature, MT-hg U, K, and T have been associated with conflicting reports, with different studies reporting either protective, risk, or non-significant effects (Ridge and Kauwe, 2018).

Alzheimer’s disease polygenic risk scores have been widely used to evaluate whether genetic liability for AD is associated with AD endophenotypes and in the prediction of disease status (Chasioti et al., 2019; Ibanez et al., 2019). These PRS, however, have generally been applied to variants across the entire genome. Using biological knowledge to incorporate variants located in genes that are part of specific pathways in the calculation of PRS, instead of considering the entire genome, allows for the construction of pathway specific PRS (Darst et al., 2017; Ibanez et al., 2019). In contrast to univariate analysis which is often underpowered due to the small effect sizes of individual SNPs, the joint analysis of the combined effect of all SNPs within a pathway may have a larger combined effect size and greater statistical power to detect an association. Pathway based analysis may also be more power predictor for understanding our specific biomarkers may contribute to disease pathogenesis. Furthermore, as a large proportion of the heritability of Alzheimer’s disease is explained by variants that lie below the genome-wide significant threshold, the inclusion of subthreshold variants allows the PRS to encompass more of the causal variance (Escott-Price et al., 2015). To date, there has been a limited application of pathway-specific PRS in AD, with only one study evaluating the association of PRSs for the immune, Aβ clearance, and cholesterol pathways with AD-related biomarkers (Darst et al., 2017; Ibanez et al., 2019). However, these PRSs were poor predictors of cognition, amyloid PET deposition, and CSF Aβ tau, and P-tau levels, potentially due to only including genome-wide significant loci (Darst et al., 2017). In the present study, we show that a pathway-specific PRS composed of SNPs located within nuclear-encoded mitochondrial genes is associated with both risk of AD and an earlier AAO, suggesting that mitochondrial function moderates AD pathogenesis. Our findings are supported by another recent study that built a molecular network using modules of coexpressed genes and identified three modules enriched for gene ontology categories related to mitochondria. These modules were associated with histopathological β-amyloid burden, cognitive decline and clinical diagnosis of AD (Mostafavi et al., 2018). Interestingly, a pathway analysis conducted by IGAP tested for overrepresentation of genes containing significantly associated SNPs within a series of functional gene sets found no evidence of enrichment in mitochondrial pathways (Jones et al., 2015). This analysis, however, only examined mitochondrial pathways that contained a subset of the mitochondrial related genes relevant to that gene set, while our study examined the aggregate effect of all nuclear encoded genes related to mitochondrial function.

The results of this study should be interpreted in conjunction with some study limitations. First, ADNI has a relatively small sample size, which can contribute to unreliable findings as a result of (a) a low probability of finding true effects, (b) a lower probability that an observed effect that is statistically significant reflects a true effect, or (c) an extracted estimate of the magnitude of an effect when a true effect is discovered (Button et al., 2013). As such the findings of this study need to be replicated in a larger cohort. Second, when constructing PRS, sample overlap between the base dataset (i.e., IGAP), and the target datasets (i.e., ADNI) can result in inflation of the association between the PRS and trait tested in the target dataset (Choi et al., 2018). However, it should be noted that IGAP consists of 54,162 participants, with ADNI only contributing 441 samples to IGAP, or 0.81% of IGAPs total sample size. Additionally, the samples included in the IGAP analysis are a subset of those included in this analysis. As such, the sample overlap between the base and target datasets is unlikely to substantially bias the results of this study. Third, the subjects in this study were of European ancestry and European MT-hg, and thus the results presented may not be generalizable to other racial/ethnic populations. In particular, in admixed populations that have a greater discordance between nuclear and mitochondrial ancestry, it could be expected that mitonuclear interactions may contribute to even more phenotypic variation in disease (Zaidi and Makova, 2019). The primary strength of this paper was evaluating the effect of MT-hgs in the context of a participant’s nuclear polygenic risk for late onset Alzheimer’s Disease (LOAD). Secondly, by imputing mtDNA variants we were able to more accurately assign MT-hgs to individuals who were included in a previous ADNI study (Lakatos et al., 2010). Finally, we utilized both cross-sectional and longitudinal data to evaluate the baseline risk of LOAD and AOO.

In conclusion, this is the first study to investigate the interactive effects of a LOAD polygenic risk score composed of mitonuclear genes and mitochondrial haplogroups on Alzheimer’s risk and survival. We found that nMT-PRS was associated with increased risk of AD and an earlier age of onset. MT-hg T was observed to attenuate the effect of the nMT-PRS on the risk of AD and AOO, while MT-hg K and V were observed to attenuate the effect of the nMT-PRS on baseline risk and strengthen the effect of the nMT-PRS on AOO respectively. The results from this study need to be replicated in independent cohorts to validate our findings. These findings suggest that interactions between the nuclear and mitochondrial genomes may influence AD pathogenesis.

## Acknowledgments

Data collection and sharing for this project was funded by the Alzheimer’s Disease Neuroimaging Initiative (ADNI) (National Institutes of Health Grant U01 AG024904) and DOD ADNI (Department of Defense award number W81XWH-12-2-0012). ADNI is funded by the National Institute on Aging, the National Institute of Biomedical Imaging and Bioengineering, and through generous contributions from the following: AbbVie, Alzheimer’s Association; Alzheimer’s Drug Discovery Foundation; Araclon Biotech; BioClinica, Inc.; Biogen; Bristol-Myers Squibb Company; CereSpir, Inc.; Cogstate; Eisai Inc.; Elan Pharmaceuticals, Inc.; Eli Lilly and Company; EuroImmun; F. Hoffmann-La Roche Ltd and its affiliated company Genentech, Inc.; Fujirebio; GE Healthcare; IXICO Ltd.; Janssen Alzheimer Immunotherapy Research & Development, LLC.; Johnson & Johnson Pharmaceutical Research & Development LLC.; Lumosity; Lundbeck; Merck & Co., Inc.; Meso Scale Diagnostics, LLC.; NeuroRx Research; Neurotrack Technologies; Novartis Pharmaceuticals Corporation; Pfizer Inc.; Piramal Imaging; Servier; Takeda Pharmaceutical Company; and Transition Therapeutics. The Canadian Institutes of Health Research is providing funds to support ADNI clinical sites in Canada. Private sector contributions are facilitated by the Foundation for the National Institutes of Health (www.fnih.org). The grantee organization is the Northern California Institute for Research and Education, and the study is coordinated by the Alzheimer’s Therapeutic Research Institute at the University of Southern California. ADNI data are disseminated by the Laboratory for Neuro Imaging at the University of Southern California.

## References

1000 Genomes Project Consortium, Auton, A., Brooks, L.D., Durbin, R.M., Garrison, E.P., Kang, H.M., Korbel, J.O., Marchini, J.L., McCarthy, S., McVean, G.A., Abecasis, G.R., 2015. A global reference for human genetic variation. Nature 526, 68–74.

Button, K.S., Ioannidis, J.P.A., Mokrysz, C., Nosek, B.A., Flint, J., Robinson, E.S.J., Munafò, M.R., 2013. Power failure: why small sample size undermines the reliability of neuroscience. Nat. Rev. Neurosci. 14, 365–376.

Calvo, S.E., Clauser, K.R., Mootha, V.K., 2016. MitoCarta2.0: an updated inventory of mammalian mitochondrial proteins. Nucleic Acids Res. 44, D1251–7.

Canevelli, M., Grande, G., Lacorte, E., Quarchioni, E., Cesari, M., Mariani, C., Bruno, G., Vanacore, N., 2016. Spontaneous Reversion of Mild Cognitive Impairment to Normal Cognition: A Systematic Review of Literature and Meta-Analysis. J. Am. Med. Dir. Assoc. 17, 943–948.

Carrieri, G., Bonafè, M., De Luca, M., Rose, G., Varcasia, O., Bruni, A., Maletta, R., Nacmias, B., Sorbi, S., Corsonello, F., Feraco, E., Andreev, K.F., Yashin, A.I., Franceschi, C., De Benedictis, G., 2001. Mitochondrial DNA haplogroups and APOE4 allele are non-independent variables in sporadic Alzheimer’s disease. Hum. Genet. 108, 194–198.

Chasioti, D., Yan, J., Nho, K., Saykin, A.J., 2019. Progress in Polygenic Composite Scores in Alzheimer’s and Other Complex Diseases. Trends Genet. 35, 371–382.

Chiba-Falek, O., Gottschalk, W.K., Lutz, M.W., 2018. The effects of the TOMM40 poly-T alleles on Alzheimer’s disease phenotypes. Alzheimers. Dement. 14, 692–698.

Chinnery, P.F., Hudson, G., 2013. Mitochondrial genetics. Br. Med. Bull. 106, 135–159.

Choi, S.W., Mak, T.S.H., O’Reilly, P., 2018. A guide to performing Polygenic Risk Score analyses. bioRxiv. https://doi.org/10.1101/416545

Coto, E., Gómez, J., Alonso, B., Corao, A.I., Díaz, M., Menéndez, M., Martínez, C., Calatayud, M.T., Morís, G., Álvarez, V., 2011. Late-onset Alzheimer’s disease is associated with mitochondrial DNA 7028C/haplogroup H and D310 poly-C tract heteroplasmy. Neurogenetics 12, 345–346.

Darst, B.F., Koscik, R.L., Racine, A.M., Oh, J.M., Krause, R.A., Carlsson, C.M., Zetterberg, H., Blennow, K., Christian, B.T., Bendlin, B.B., Okonkwo, O.C., Hogan, K.J., Hermann, B.P., Sager, M.A., Asthana, S., Johnson, S.C., Engelman, C.D., 2017. Pathway-Specific Polygenic Risk Scores as Predictors of Amyloid-β-Deposition and Cognitive Function in a Sample at Increased Risk for Alzheimer’s Disease. J. Alzheimers. Dis. 55, 473–484.

de Leeuw, C.A., Mooij, J.M., Heskes, T., Posthuma, D., 2015. MAGMA: generalized gene-set analysis of GWAS data. PLoS Comput. Biol. 11, e1004219.

Dobler, R., Dowling, D.K., Morrow, E.H., Reinhardt, K., 2018. A systematic review and meta-analysis reveals pervasive effects of germline mitochondrial replacement on components of health. Hum. Reprod. Update 24, 519–534.

Edland, S.D., Tobe, V.O., Rieder, M.J., Bowen, J.D., McCormick, W., Teri, L., Schellenberg, G.D., Larson, E.B., Nickerson, D.A., Kukull, W.A., 2002. Mitochondrial genetic variants and Alzheimer disease: a case-control study of the T4336C and G5460A variants. Alzheimer Dis. Assoc. Disord. 16, 1–7.

Escott-Price, V., Sims, R., Bannister, C., Harold, D., Vronskaya, M., Majounie, E., Badarinarayan, N., GERAD/PERADES, IGAP consortia, Morgan, K., Passmore, P., Holmes, C., Powell, J., Brayne, C., Gill, M., Mead, S., Goate, A., Cruchaga, C., Lambert, J.-C., van Duijn, C., Maier, W., Ramirez, A., Holmans, P., Jones, L., Hardy, J., Seshadri, S., Schellenberg, G.D., Amouyel, P., Williams, J., 2015. Common polygenic variation enhances risk prediction for Alzheimer’s disease. Brain 138, 3673–3684.

Euesden, J., Lewis, C.M., O’Reilly, P.F., 2015. PRSice: Polygenic Risk Score software. Bioinformatics 31, 1466–1468.

Gorman, G.S., Chinnery, P.F., DiMauro, S., Hirano, M., Koga, Y., McFarland, R., Suomalainen, A., Thorburn, D.R., Zeviani, M., Turnbull, D.M., 2016. Mitochondrial diseases. Nat Rev Dis Primers 2, 16080.

Honea, R.A., Vidoni, E.D., Swerdlow, R.H., Burns, J.M., Alzheimer’s Disease Neuroimaging Initiative, 2012. Maternal family history is associated with Alzheimer’s disease biomarkers. J. Alzheimers. Dis. 31, 659–668.

Howie, B.N., Donnelly, P., Marchini, J., 2009. A flexible and accurate genotype imputation method for the next generation of genome-wide association studies. PLoS Genet. 5, e1000529.

Ibanez, L., Farias, F.H.G., Dube, U., Mihindukulasuriya, K.A., Harari, O., 2019. Polygenic Risk Scores in Neurodegenerative Diseases: a Review. Curr. Genet. Med. Rep. 7, 22–29.

Jones, L., Lambert, J.-C., Wang, L.-S., Choi, S.-H., Harold, D., Vedernikov, A., Escott-Price, V., Stone, T., Richards, A., Bellenguez, C., Ibrahim-Verbaas, C.A., Naj, A.C., Sims, R., Gerrish, A., Jun, G., DeStefano, A.L., Bis, J.C., Beecham, G.W., Grenier-Boley, B., Russo, G., Thornton-Wells, T.A., Jones, N., Smith, A.V., Chouraki, V., Thomas, C., Ikram, M.A., Zelenika, D., Vardarajan, B.N., Kamatani, Y., Lin, C.-F., Schmidt, H., Kunkle, B.W., Dunstan, M.L., Ruiz, A., Bihoreau, M.-T., Reitz, C., Pasquier, F., Hollingworth, P., Hanon, O., Fitzpatrick, A.L., Buxbaum, J., Campion, D., Crane, P.K., Becker, T., Gudnason, V., Cruchaga, C., Craig, D., Amin, N., Berr, C., Lopez, O.L., De Jager, P.L., Deramecourt, V., Johnston, J.A., Evans, D., Lovestone, S., Letteneur, L., Kornhuber, J., Tárraga, L., Rubinsztein, D.C., Eiriksdottir, G., Sleegers, K., Goate, A.M., Fiévet, N., Huentelman, M.J., Gill, M., Emilsson, V., Brown, K., Kamboh, M.I., Keller, L., Barberger-Gateau, P., McGuinness, B., Larson, E.B., Myers, A.J., Dufouil, C., Todd, S., Wallon, D., Love, S., Kehoe, P., Rogaeva, E., Gallacher, J., St George-Hyslop, P., Clarimon, J., Lleò, A., Bayer, A., Tsuang, D.W., Yu, L., Tsolaki, M., Bossù, P., Spalletta, G., Proitsi, P., Collinge, J., Sorbi, S., Sanchez Garcia, F., Fox, N., Hardy, J., Deniz Naranjo, M.C., Razquin, C., Bosco, P., Clarke, R., Brayne, C., Galimberti, D., Mancuso, M., Moebus, S., Mecocci, P., del Zompo, M., Maier, W., Hampel, H., Pilotto, A., Bullido, M., Panza, F., Caffarra, P., Nacmias, B., Gilbert, J.R., Mayhaus, M., Jessen, F., Dichgans, M., Lannfelt, L., Hakonarson, H., Pichler, S., Carrasquillo, M.M., Ingelsson, M., Beekly, D., Alavarez, V., Zou, F., Valladares, O., Younkin, S.G., Coto, E., Hamilton-Nelson, K.L., Mateo, I., Owen, M.J., Faber, K.M., Jonsson, P.V., Combarros, O., O’Donovan, M.C., Cantwell, L.B., Soininen, H., Blacker, D., Mead, S., Mosley, T.H., Bennett, D.A., Harris, T.B., Fratiglioni, L., Holmes, C., de Bruijn, R.F.A.G., Passmore, P., Montine, T.J., Bettens, K., Rotter, J.I., Brice, A., Morgan, K., Foroud, T.M., Kukull, W.A., Hannequin, D., Powell, J.F., Nalls, M.A., Ritchie, K., Lunetta, K.L., Kauwe, J.S.K., Boerwinkle, E., Riemenschneider, M., Boada, M., Hiltunen, M., Martin, E.R., Pastor, P., Schmidt, R., Rujescu, D., Dartigues, J.-F., Mayeux, R., Tzourio, C., Hofman, A., Nöthen, M.M., Graff, C., Psaty, B.M., Haines, J.L., Lathrop, M., Pericak-Vance, M.A., Launer, L.J., Farrer, L.A., van Duijn, C.M., Van Broeckhoven, C., Ramirez, A., Schellenberg, G.D., Seshadri, S., Amouyel, P., Williams, J., Holmans, P.A., 2015. Convergent genetic and expression data implicate immunity in Alzheimer’s disease. Alzheimers. Dement. 11, 658–671.

Kokotas, H., Petersen, M.B., Willems, P.J., 2007. Mitochondrial deafness. Clin. Genet. 71, 379–391.

Kunkle, B.W., Grenier-Boley, B., Sims, R., Bis, J.C., Damotte, V., Naj, A.C., Boland, A., Vronskaya, M., van der Lee, S.J., Amlie-Wolf, A., Bellenguez, C., Frizatti, A., Chouraki, V., Martin, E.R., Sleegers, K., Badarinarayan, N., Jakobsdottir, J., Hamilton-Nelson, K.L., Moreno-Grau, S., Olaso, R., Raybould, R., Chen, Y., Kuzma, A.B., Hiltunen, M., Morgan, T., Ahmad, S., Vardarajan, B.N., Epelbaum, J., Hoffmann, P., Boada, M., Beecham, G.W., Garnier, J.-G., Harold, D., Fitzpatrick, A.L., Valladares, O., Moutet, M.-L., Gerrish, A., Smith, A.V., Qu, L., Bacq, D., Denning, N., Jian, X., Zhao, Y., Del Zompo, M., Fox, N.C., Choi, S.-H., Mateo, I., Hughes, J.T., Adams, H.H., Malamon, J., Sanchez-Garcia, F., Patel, Y., Brody, J.A., Dombroski, B.A., Naranjo, M.C.D., Daniilidou, M., Eiriksdottir, G., Mukherjee, S., Wallon, D., Uphill, J., Aspelund, T., Cantwell, L.B., Garzia, F., Galimberti, D., Hofer, E., Butkiewicz, M., Fin, B., Scarpini, E., Sarnowski, C., Bush, W.S., Meslage, S., Kornhuber, J., White, C.C., Song, Y., Barber, R.C., Engelborghs, S., Sordon, S., Voijnovic, D., Adams, P.M., Vandenberghe, R., Mayhaus, M., Cupples, L.A., Albert, M.S., De Deyn, P.P., Gu, W., Himali, J.J., Beekly, D., Squassina, A., Hartmann, A.M., Orellana, A., Blacker, D., Rodriguez-Rodriguez, E., Lovestone, S., Garcia, M.E., Doody, R.S., Munoz-Fernadez, C., Sussams, R., Lin, H., Fairchild, T.J., Benito, Y.A., Holmes, C., Karamujić-Ćomić H., Frosch, M.P., Thonberg, H., Maier, W., Roschupkin, G., Ghetti, B., Giedraitis, V., Kawalia, A., Li, S., Huebinger, R.M., Kilander, L., Moebus, S., Hernández, I., Kamboh, M.I., Brundin, R., Turton, J., Yang, Q., Katz, M.J., Concari, L., Lord, J., Beiser, A.S., Keene, C.D., Helisalmi, S., Kloszewska, I., Kukull, W.A., Koivisto, A.M., Lynch, A., Tarraga, L., Larson, E.B., Haapasalo, A., Lawlor, B., Mosley, T.H., Lipton, R.B., Solfrizzi, V., Gill, M., Longstreth, W.T., Montine, T.J., Frisardi, V., Diez-Fairen, M., Rivadeneira, F., Petersen, R.C., Deramecourt, V., Alvarez, I., Salani, F., Ciaramella, A., Boerwinkle, E., Reiman, E.M., Fievet, N., Rotter, J.I., Reisch, J.S., Hanon, O., Cupidi, C., Andre Uitterlinden, A.G., Royall, D.R., Dufouil, C., Maletta, R.G., de Rojas, I., Sano, M., Brice, A., Cecchetti, R., George-Hyslop, P.S., Ritchie, K., Tsolaki, M., Tsuang, D.W., Dubois, B., Craig, D., Wu, C.-K., Soininen, H., Avramidou, D., Albin, R.L., Fratiglioni, L., Germanou, A., Apostolova, L.G., Keller, L., Koutroumani, M., Arnold, S.E., Panza, F., Gkatzima, O., Asthana, S., Hannequin, D., Whitehead, P., Atwood, C.S., Caffarra, P., Hampel, H., Quintela, I., Carracedo, Á., Lannfelt, L., Rubinsztein, D.C., Barnes, L.L., Pasquier, F., Frölich, L., Barral, S., McGuinness, B., Beach, T.G., Johnston, J.A., Becker, J.T., Passmore, P., Bigio, E.H., Schott, J.M., Bird, T.D., Warren, J.D., Boeve, B.F., Lupton, M.K., Bowen, J.D., Proitsi, P., Boxer, A., Powell, J.F., Burke, J.R., Kauwe, J.S.K., Burns, J.M., Mancuso, M., Buxbaum, J., Bonuccelli, U., Cairns, N.J., McQuillin, A., Cao, C., Livingston, G., Carlson, C.S., Bass, N.J., Carlsson, C.M., Hardy, J., Carney, R.M., Bras, J., Carrasquillo, M.M., Guerreiro, R., Allen, M., Chui, H.C., Fisher, E., Masullo, C., Crocco, E.A., DeCarli, C., Bisceglio, G., Dick, M., Ma, L., Duara, R., Graff-Radford, N.R., Evans, D.A., Hodges, A., Faber, K.M., Scherer, M., Fallon, K.B., Riemenschneider, M., Fardo, D.W., Heun, R., Farlow, M.R., Kölsch, H., Ferris, S., Leber, M., Foroud, T.M., Heuser, I., Galasko, D.R., Giegling, I., Gearing, M., Hüll, M., Geschwind, D.H., Gilbert, J.R., Morris, J., Green, R.C., Mayo, K., Growdon, J.H., Feulner, T., Hamilton, R.L., Harrell, L.E., Drichel, D., Honig, L.S., Cushion, T.D., Huentelman, M.J., Hollingworth, P., Hulette, C.M., Hyman, B.T., Marshall, R., Jarvik, G.P., Meggy, A., Abner, E., Menzies, G.E., Jin, L.-W., Leonenko, G., Real, L.M., Jun, G.R., Baldwin, C.T., Grozeva, D., Karydas, A., Russo, G., Kaye, J.A., Kim, R., Jessen, F., Kowall, N.W., Vellas, B., Kramer, J.H., Vardy, E., LaFerla, F.M., Jöckel, K.-H., Lah, J.J., Dichgans, M., Leverenz, J.B., Mann, D., Levey, A.I., Pickering-Brown, S., Lieberman, A.P., 2019. Genetic meta-analysis of diagnosed Alzheimer’s disease identifies new risk loci and implicates Aβ, tau, immunity and lipid processing. Nat. Genet. https://doi.org/10.1038/s41588-019-0358-2

Lakatos, A., Derbeneva, O., Younes, D., Keator, D., Bakken, T., Lvova, M., Brandon, M., Guffanti, G., Reglodi, D., Saykin, A., Weiner, M., Macciardi, F., Schork, N., Wallace, D.C., Potkin, S.G., Alzheimer’s Disease Neuroimaging Initiative, 2010. Association between mitochondrial DNA variations and Alzheimer’s disease in the ADNI cohort. Neurobiol. Aging 31, 1355–1363.

Lambert, J.C., Ibrahim-Verbaas, C.A., Harold, D., Naj, A.C., Sims, R., Bellenguez, C., DeStafano, A.L., Bis, J.C., Beecham, G.W., Grenier-Boley, B., Russo, G., Thorton-Wells, T.A., Jones, N., Smith, A.V., Chouraki, V., Thomas, C., Ikram, M.A., Zelenika, D., Vardarajan, B.N., Kamatani, Y., Lin, C.F., Gerrish, A., Schmidt, H., Kunkle, B., Dunstan, M.L., Ruiz, A., Bihoreau, M.T., Choi, S.H., Reitz, C., Pasquier, F., Cruchaga, C., Craig, D., Amin, N., Berr, C., Lopez, O.L., De Jager, P.L., Deramecourt, V., Johnston, J.A., Evans, D., Lovestone, S., Letenneur, L., Morón, F.J., Rubinsztein, D.C., Eiriksdottir, G., Sleegers, K., Goate, A.M., Fiévet, N., Huentelman, M.W., Gill, M., Brown, K., Kamboh, M.I., Keller, L., Barberger-Gateau, P., McGuiness, B., Larson, E.B., Green, R., Myers, A.J., Dufouil, C., Todd, S., Wallon, D., Love, S., Rogaeva, E., Gallacher, J., St George-Hyslop, P., Clarimon, J., Lleo, A., Bayer, A., Tsuang, D.W., Yu, L., Tsolaki, M., Bossù, P., Spalletta, G., Proitsi, P., Collinge, J., Sorbi, S., Sanchez-Garcia, F., Fox, N.C., Hardy, J., Deniz Naranjo, M.C., Bosco, P., Clarke, R., Brayne, C., Galimberti, D., Mancuso, M., Matthews, F., European Alzheimer’s Disease Initiative (EADI), Genetic and Environmental Risk in Alzheimer’s Disease, Alzheimer’s Disease Genetic Consortium, Cohorts for Heart and Aging Research in Genomic Epidemiology, Moebus, S., Mecocci, P., Del Zompo, M., Maier, W., Hampel, H., Pilotto, A., Bullido, M., Panza, F., Caffarra, P., Nacmias, B., Gilbert, J.R., Mayhaus, M., Lannefelt, L., Hakonarson, H., Pichler, S., Carrasquillo, M.M., Ingelsson, M., Beekly, D., Alvarez, V., Zou, F., Valladares, O., Younkin, S.G., Coto, E., Hamilton-Nelson, K.L., Gu, W., Razquin, C., Pastor, P., Mateo, I., Owen, M.J., Faber, K.M., Jonsson, P.V., Combarros, O., O’Donovan, M.C., Cantwell, L.B., Soininen, H., Blacker, D., Mead, S., Mosley, T.H., Jr, Bennett, D.A., Harris, T.B., Fratiglioni, L., Holmes, C., de Bruijn, R.F., Passmore, P., Montine, T.J., Bettens, K., Rotter, J.I., Brice, A., Morgan, K., Foroud, T.M., Kukull, W.A., Hannequin, D., Powell, J.F., Nalls, M.A., Ritchie, K., Lunetta, K.L., Kauwe, J.S., Boerwinkle, E., Riemenschneider, M., Boada, M., Hiltuenen, M., Martin, E.R., Schmidt, R., Rujescu, D., Wang, L.S., Dartigues, J.F., Mayeux, R., Tzourio, C., Hofman, A., Nöthen, M.M., Graff, C., Psaty, B.M., Jones, L., Haines, J.L., Holmans, P.A., Lathrop, M., Pericak-Vance, M.A., Launer, L.J., Farrer, L.A., van Duijn, C.M., Van Broeckhoven, C., Moskvina, V., Seshadri, S., Williams, J., Schellenberg, G.D., Amouyel, P., 2013. Meta-analysis of 74,046 individuals identifies 11 new susceptibility loci for Alzheimer’s disease. Nat. Genet. 45, 1452–1458.

Lehner, B., 2011. Molecular mechanisms of epistasis within and between genes. Trends Genet. 27, 323–331.

Maruszak, A., Safranow, K., Branicki, W., Gawęda-Walerych, K., Pośpiech, E., Gabryelewicz, T., Canter, J.A., Barcikowska, M., Zekanowski, C., 2011. The impact of mitochondrial and nuclear DNA variants on late-onset Alzheimer’s disease risk. J. Alzheimers. Dis. 27, 197–210.

Masters, C.L., Bateman, R., Blennow, K., Rowe, C.C., Sperling, R.A., Cummings, J.L., 2015. Alzheimer’s disease. Nat Rev Dis Primers 1, 15056.

McCarthy, S., Das, S., Kretzschmar, W., Delaneau, O., Wood, A.R., Teumer, A., Kang, H.M., Fuchsberger, C., Danecek, P., Sharp, K., Luo, Y., Sidore, C., Kwong, A., Timpson, N., Koskinen, S., Vrieze, S., Scott, L.J., Zhang, H., Mahajan, A., Veldink, J., Peters, U., Pato, C., van Duijn, C.M., Gillies, C.E., Gandin, I., Mezzavilla, M., Gilly, A., Cocca, M., Traglia, M., Angius, A., Barrett, J.C., Boomsma, D., Branham, K., Breen, G., Brummett, C.M., Busonero, F., Campbell, H., Chan, A., Chen, S., Chew, E., Collins, F.S., Corbin, L.J., Smith, G.D., Dedoussis, G., Dorr, M., Farmaki, A.-E., Ferrucci, L., Forer, L., Fraser, R.M., Gabriel, S., Levy, S., Groop, L., Harrison, T., Hattersley, A., Holmen, O.L., Hveem, K., Kretzler, M., Lee, J.C., McGue, M., Meitinger, T., Melzer, D., Min, J.L., Mohlke, K.L., Vincent, J.B., Nauck, M., Nickerson, D., Palotie, A., Pato, M., Pirastu, N., McInnis, M., Richards, J.B., Sala, C., Salomaa, V., Schlessinger, D., Schoenherr, S., Slagboom, P.E., Small, K., Spector, T., Stambolian, D., Tuke, M., Tuomilehto, J., Van den Berg, L.H., Van Rheenen, W., Volker, U., Wijmenga, C., Toniolo, D., Zeggini, E., Gasparini, P., Sampson, M.G., Wilson, J.F., Frayling, T., de Bakker, P.I.W., Swertz, M.A., McCarroll, S., Kooperberg, C., Dekker, A., Altshuler, D., Willer, C., Iacono, W., Ripatti, S., Soranzo, N., Walter, K., Swaroop, A., Cucca, F., Anderson, C.A., Myers, R.M., Boehnke, M., McCarthy, M.I., Durbin, R., Haplotype Reference Consortium, 2016. A reference panel of 64,976 haplotypes for genotype imputation. Nat. Genet. 48, 1279–1283.

McInerney, T.W., Fulton-Howard, B., Patterson, C., Paliwal, D., Jermiin, L.S., Patel, H., Pa, J., Swerdlow, R.H., Goate, A., Easteal, S., Andrews, S.J., for the Alzheimer’s Disease Neuroimaging Initiative, 2019. MitoImpute: A Snakemake pipeline for imputation of mitochondrial genetic variants. bioRxiv. https://doi.org/10.1101/649293

Mhatre, S.D., Tsai, C.A., Rubin, A.J., James, M.L., Andreasson, K.I., 2015. Microglial malfunction: the third rail in the development of Alzheimer’s disease. Trends Neurosci. 38, 621–636.

Morrow, E.H., Camus, M.F., 2017. Mitonuclear epistasis and mitochondrial disease. Mitochondrion 35, 119–122.

Mostafavi, S., Gaiteri, C., Sullivan, S.E., White, C.C., Tasaki, S., Xu, J., Taga, M., Klein, H.-U., Patrick, E., Komashko, V., McCabe, C., Smith, R., Bradshaw, E.M., Root, D.E., Regev, A., Yu, L., Chibnik, L.B., Schneider, J.A., Young-Pearse, T.L., Bennett, D.A., De Jager, P.L., 2018. A molecular network of the aging human brain provides insights into the pathology and cognitive decline of Alzheimer’s disease. Nat. Neurosci. 21, 811–819.

Navarro-Gomez, D., Leipzig, J., Shen, L., Lott, M., Stassen, A.P.M., Wallace, D.C., Wiggs, J.L., Falk, M.J., van Oven, M., Gai, X., 2015. Phy-Mer: a novel alignment-free and reference-independent mitochondrial haplogroup classifier. Bioinformatics 31, 1310–1312.

Panza, F., Lozupone, M., Logroscino, G., Imbimbo, B.P., 2019. A critical appraisal of amyloid-β-targeting therapies for Alzheimer disease. Nat. Rev. Neurol. https://doi.org/10.1038/s41582-018-0116-6

Perez Ortiz, J.M., Swerdlow, R.H., 2019. Mitochondrial dysfunction in Alzheimer’s disease: Role in pathogenesis and novel therapeutic opportunities. Br. J. Pharmacol. https://doi.org/10.1111/bph.14585

Purcell, S., Neale, B., Todd-Brown, K., Thomas, L., Ferreira, M.A.R., Bender, D., Maller, J., Sklar, P., de Bakker, P.I.W., Daly, M.J., Sham, P.C., 2007. PLINK: a tool set for whole-genome association and population-based linkage analyses. Am. J. Hum. Genet. 81, 559–575.

Ridge, P.G., Kauwe, J.S.K., 2018. Mitochondria and Alzheimer’s Disease: the Role of Mitochondrial Genetic Variation. Curr. Genet. Med. Rep. 6, 1–10.

Ridge, P.G., Koop, A., Maxwell, T.J., Bailey, M.H., Swerdlow, R.H., Kauwe, J.S.K., Honea, R.A., Alzheimer’s Disease Neuroimaging Initiative, 2013. Mitochondrial haplotypes associated with biomarkers for Alzheimer’s disease. PLoS One 8, e74158.

Ridge, P.G., Wadsworth, M.E., Miller, J.B., Saykin, A.J., Green, R.C., Alzheimer’s Disease Neuroimaging Initiative, Kauwe, J.S.K., 2018. Assembly of 809 whole mitochondrial genomes with clinical, imaging, and fluid biomarker phenotyping. Alzheimers. Dement. 14, 514–519.

Saykin, A.J., Shen, L., Yao, X., Kim, S., Nho, K., Risacher, S.L., Ramanan, V.K., Foroud, T.M., Faber, K.M., Sarwar, N., Munsie, L.M., Hu, X., Soares, H.D., Potkin, S.G., Thompson, P.M., Kauwe, J.S.K., Kaddurah-Daouk, R., Green, R.C., Toga, A.W., Weiner, M.W., Alzheimer’s Disease Neuroimaging Initiative, 2015. Genetic studies of quantitative MCI and AD phenotypes in ADNI: Progress, opportunities, and plans. Alzheimers. Dement. 11, 792–814.

Swerdlow, R.H., 2018. Mitochondria and Mitochondrial Cascades in Alzheimer’s Disease. J. Alzheimers. Dis. 62, 1403–1416.

Taanman, J.W., 1999. The mitochondrial genome: structure, transcription, translation and replication. Biochim. Biophys. Acta 1410, 103–123.

van Oven, M., Kayser, M., 2009. Updated comprehensive phylogenetic tree of global human mitochondrial DNA variation. Hum. Mutat. 30, E386–94.

Weissensteiner, H., Pacher, D., Kloss-Brandstätter, A., Forer, L., Specht, G., Bandelt, H.-J., Kronenberg, F., Salas, A., Schönherr, S., 2016. HaploGrep 2: mitochondrial haplogroup classification in the era of high-throughput sequencing. Nucleic Acids Res. 44, W58–63.

Zaidi, A.A., Makova, K.D., 2019. Investigating mitonuclear interactions in human admixed populations. Nat Ecol Evol 3, 213–222.

