## Supplementary material for "Mitonuclear interactions influence Alzheimer’s disease risk": Table S1

Supplementary Table 1: Association of mitonuclear interactions with baseline risk of Alzheimer's disease and Alzheimer’s age of onset adjusting for PRS excluding nMT genes and APOE.

|  | Cross-sectional | | Survival | |
| --- | --- | --- | --- | --- |
| Variable | Main | Interaction | Main | Interaction |
| Age | 0 (0.02) | 0.01 (0.02) | - | - |
| Male | 0.16 (0.28) | 0.03 (0.29) | -0.34 (0.1)*** | -0.35 (0.1)*** |
| APOE status |  |  |  |  |
| ε4+ | 1.66 (0.32)*** | 1.72 (0.33)*** | 0.78 (0.11)*** | 0.77 (0.12)*** |
| ε2+ | -0.6 (0.72) | -0.74 (0.74) | -0.71 (0.39). | -0.73 (0.39). |
| PC1 | -0.59 (0.17)*** | -0.55 (0.17)*** | -0.3 (0.07)*** | -0.28 (0.07)*** |
| PC2 | 0.84 (0.69) | 0.69 (0.7) | 0.05 (0.04) | 0.05 (0.04) |
| PRS w/o nMT & APOE | 3.26 (0.33)*** | 3.26 (0.34)*** | 1.03 (0.09)*** | 1.03 (0.09)*** |
| nMT-PRS | 0.42 (0.17)* | 0.68 (0.24)** | 0.15 (0.06)* | 0.17 (0.08)* |
| Haplogroup |  |  |  |  |
| I | 0.55 (0.74) | 0.63 (0.74) | -0.26 (0.26) | -0.15 (0.28) |
| J | 0.53 (0.55) | 0.55 (0.57) | 0.08 (0.18) | 0.06 (0.19) |
| K | 0.48 (0.45) | 0.44 (0.44) | -0.11 (0.18) | -0.09 (0.19) |
| T | 0.43 (0.47) | 0.56 (0.48) | -0.14 (0.17) | -0.08 (0.18) |
| U | 1.01 (0.46)* | 1.11 (0.48)* | -0.13 (0.16) | -0.14 (0.17) |
| V | 0.5 (0.8) | 0.81 (1.02) | 0.22 (0.28) | -0.29 (0.41) |
| W | 0.19 (1.14) | 0.55 (1.14) | 0.47 (0.33) | 0.5 (0.33) |
| X | -0.29 (1.23) | -1.51 (2.48) | -0.02 (0.33) | 0.21 (0.37) |
| Haplogroup x nMT-PRS | |  |  |  |
| I | - | -0.36 (1.15) | - | -0.26 (0.28) |
| J | - | 0.11 (0.8) | - | 0.09 (0.22) |
| K | - | -0.9 (0.46). | - | -0.11 (0.17) |
| T | - | -1.25 (0.58)* | - | -0.22 (0.21) |
| U | - | 0.24 (0.61) | - | 0.01 (0.16) |
| V | - | 1.16 (1.93) | - | 0.88 (0.39)* |
| W | - | -1.02 (0.99) | - | 0.26 (0.35) |
| X | - | -3.29 (2.78) | - | -0.37 (0.34) |

‘.’ p < 0.1; * p < 0.05; ** p < 0.01; *** p < 0.001
